# High Throughput Transcriptomics to Understand Chemical Drivers of Aggressive Breast Cancer Subtypes

**DOI:** 10.1101/2022.11.16.516817

**Authors:** Kimberley E. Sala-Hamrick, Anagha Tapaswi, Katelyn M. Polemi, Vy. K Nguyen, Justin A. Colacino

## Abstract

**Background:** The impact of chemical exposures on breast cancer progression is poorly characterized and may influence the development of more severe and aggressive subtypes.

**Objectives:** There is a suite of toxicants, including metals, pesticides, and personal care product compounds, which are commonly detected at high levels in US Center for Disease Control’s National Health and Nutrition Examination Survey (NHANES) chemical biomarker screens. To characterize the impact of these toxicants on breast cancer pathways, we performed high throughput dose-response transcriptomic analysis of toxicant exposed breast cells.

**Methods:** We treated non-tumorigenic mammary epithelial cells, MCF10A, with 21 chemicals at four doses (25nM, 250nM, 2.5µM, 25µM) for 48 hours. We conducted RNA-sequencing for these 408 samples, adapting the PlexWell plate-based RNA-sequencing method to analyze changes in gene expression resulting from these exposures. For each chemical, we calculated gene and biological pathway specific benchmark doses using BMDExpress2, identifying differentially expressed genes and generating the best fit benchmark dose models for each gene. We employed enrichment testing to test whether each chemical’s upregulated or downregulated genes were over-represented in a biological process or pathway. We contextualized benchmark doses relative to human population biomarker concentrations in NHANES.

**Results:** Overall, significant changes in gene expression varied across doses of each chemical and benchmark dose modeling revealed dose-responsive alterations of thousands of different genes. Comparison of benchmark data to NHANES chemical biomarker concentrations indicated an overlap between actual exposure levels and levels sufficient to cause a gene expression response. Enrichment and cell deconvolution analyses showed benchmark dose responses correlated with changes in cancer and breast cancer related pathways, including induction of basal-like characteristics for some chemicals, including p,p’-DDE, lead, copper, and methyl paraben.

**Discussion:** These analyses revealed that these 21 chemicals induce significant changes in pathways involved in breast cancer initiation and progression at human exposure relevant doses.

## Introduction

There are an estimated 2.3 million new cases of breast cancer diagnosed globally and approximately 685,000 people die from breast cancer each year .^1^ Breast cancer is a heterogeneous disease – tumors can be grouped into subtypes based on their molecular signature and expression of estrogen receptor (ER), progesterone receptor (PR), and human epidermal growth factor receptor 2 (HER2).^2^ Prognosis and outcomes vary based on subtype. Triple negative breast cancers, those which do not express ER, PR, or HER2, have the worst overall survival rates.^3^ Individuals diagnosed with metastatic breast cancer continue to have very poor survival, less than 30% after five years, regardless of subtype.^4^ Identifying the factors which promote these aggressive breast cancers, such as metastatic and/or triple negative breast cancers, is of high public health priority. In general, high penetrance genetic risk factors are only thought to explain approximately 5-10% of all breast cancers.^5^ As such, there is likely a large impact of the environment on aggressive breast cancer risk. There is an urgent need to identify potentially modifiable risk factors, such as chemical exposures, which promote aggressive breast cancers so that we can develop new strategies for prevention and targeted therapies.

As aggressive breast cancers form, they acquire the Hallmarks of Cancer, including sustained proliferative signaling, the activation of invasion and metastasis pathways, immortality, and the evasion of growth suppression, the reactivation of stem cell pathways, and the acquisition of cellular plasticity.^6–8^ As an example, triple negative breast cancers and breast cancers which are more likely to metastasize are enriched for expression of stem cell associated pathways.^9-13^ Triple negative breast cancers also often present with a basal-like phenotype.^14^ Experimental data surprisingly suggest that these basal-like breast cancers develop from dysregulated luminal cells, where these cells acquire phenotypic plasticity and shift from a luminal-like to a basal-like expression pattern.^15,16^ Identifying chemicals which impact the development of aggressive breast cancers through the promotion of these various hallmarks, including altered stemness and phenotypic plasticity pathways, would provide novel insights into the mechanisms by which environmental exposures impact breast cancer risk and outcomes.

There are hundreds or thousands of chemical exposures which could impact breast cancer risk, and a systematic and rapid method for evaluation is essential to identify those which may alter aggressive breast cancer hallmarks at human relevant doses.^5^ Here, we developed a high throughput transcriptomic platform for the unbiased assessment of chemical effects *in vitro*. We quantified the dose-dependent effects of 21 chemical exposures in the commonly used *in vitro* breast carcinogenesis model, non-tumorigenic MCF10A cells. We prioritized the chemicals based on them being commonly detected in chemical biomarker screens, with documented exposure disparities across demographic groups, and with putative links to breast cancer.^5, 17-23^ Across over 400 RNA-sequencing reactions, we characterize the dose-response effects of these chemicals on individual genes, breast cancer hallmarks, and on cellular plasticity phenotypes. We contextualize our dose-response data with respect to detectable biomarker concentrations in a representative sample of the US population. These results highlight putative chemical contributors to aggressive breast cancers which occur at human relevant doses and provide a methodological framework for the evaluation of chemical exposures on biological pathways associated with aggressive breast cancers.

## Methods

### Cell Culture

The non-tumorigenic mammary epithelial cell line, MCF10A, was obtained from American Type Culture Collection (ATCC, Manassas, VA, USA). Cells were cultured and maintained in Dulbecco’s Modified Eagle’s Medium/Hams F12 50/50 mix (Corning, Corning, NY, USA) supplemented with 5% Horse Serum (Thermo Fisher, Waltham, MA, USA), 2.5mg/mL HEPES (Thermo Fisher, Waltham, MA, USA), 5ug/mL Insulin (Gibco/Thermo Fisher, Waltham, MA, USA), 96ug/mL Hydrocortisone (StemCell, Vancouver, Canada), 100ug/mL Cholera toxin (Sigma-Aldrich, Saint Louis, MO, USA) and 20ng/mL Epidermal Growth Factor (StemCell, Vancouver, Canada). The cells were maintained at 37^°^C in a humidified 5% CO_2_ condition.

### Compound preparation

The 21 chemicals were purchased from Sigma-Aldrich (Saint Louis, MO, USA) and Cayman Chemicals (Ann Arbor, MI, USA) (Supplementary Table 1). The chemicals were weighed and dissolved in Dimethyl sulfoxide (DMSO) (for most) and in Sterile Water (for some) at a concentration of 5mg/mL and stored at -20^°^C for long term storage. For dosing, we prepared an intermediate stock concentration of 5mM for each of the chemicals, which was further diluted into final concentrations for dosing the MCF10A cells: 25uM, 2.5uM, 0.25uM and 0.025uM. These final concentrations were prepared fresh in MCF10A media for each experiment. For chemicals dissolved in DMSO, the final DMSO concentration was 0.5%, as were the vehicle controls. For chemicals dissolved in water, an equivalent amount of water was included in the vehicle control.

### Cell plating

MCF10A cells were cultured and expanded at passage 105 for all rounds of cell culture. Cells were plated in black, clear flat bottom 384 well plates (Corning, Corning, NY, USA) using a 2.5-125uL multi-channel pipette. First, 10uL of plain media was added to pre-warmed plates. 500 cells in 30 uL of media were added to each well at a slow speed to make up a final volume of 40 uL. The cells were left undisturbed to allow the cells to attach and distribute uniformly over the well. Each microplate was incubated at 37^°^C humidified incubator with 5% CO_2_ for 24 hours prior to dosing. Cells were then exposed to test chemicals for 48 hours with 4 biological replicates per dose. Experiments were run in 2 batches to include experimental replicates for all chemical-dose combinations.

### plexWell RNA-Sequencing Overview

To conduct high throughput RNA-sequencing, we developed an adapted version of the plexWell (SeqWell, Beverly, MA, USA) plate-based sequencing method to make the method applicable for high throughput *in vitro* toxicological analysis. In the plexWell protocol, sequencing reactions are prepared in custom 96 well plates, where each well contains a unique oligo barcode. Plates are also assigned a unique oligo barcode. Cells are lysed and cDNA is directly prepared from the cellular lysate. A tagmentation reaction adds the well specific barcode to the cDNA from each well. The tagmented cDNAs are then pooled prior to library preparation, where plate specific barcodes can also be incorporated for ultrahigh sample multiplexing prior to sequencing. For example, 384 wells of treated samples would be processed across 4 custom 96 well plates, where each of the samples receives a well specific barcode and each of the 4 plates receives a plate specific barcode. This leads to each of the 384 samples receiving a unique combination of well and plate barcodes, which can be bioinformatically deconvoluted following sequencing. This protocol decreases the cost of library preparation by approximately 90% relative to traditional RNA-sequencing, enabling unbiased high throughput toxicological analyses.

Our key modification to the manufacturer protocol to streamline the process was to avoid the step of RNA purification prior to cDNA preparation. The plexWell protocol calls for the input of 1uL of RNA for cDNA preparation. To eliminate the step of RNA extraction, we devised a method of lysing the cells directly in the 384-well microplate. After 48 hours of incubation with the test chemicals, the cells were washed twice with 40uL of 1X HBSS. After the 2nd wash, 5uL of 1X HBSS was left behind, to avoid cell loss. We added 10uL of plexWell lysis buffer (with RNase inhibitor) to each well using a 2.5-125uL multi-channel micropipette. The buffer was pipetted up and down to mix 10 times and the cell lysates were collected in 1mL eppendorf tubes on ice to avoid RNA degradation. Cell lysis was confirmed via examination by microscopy.

### plexWell cDNA preparation

The plexWell manufacturer’s protocol was followed to prepare cDNA for subsequent library preparation. In brief, 1uL of cell lysate was used as an input for cDNA preparation and oligo dT annealing in a PCR 96 well plate (Dot Scientific, Burton, MI, USA). The cDNA synthesis reaction was run for 12 cycles on a Bio-Rad CFX96 Touch Real-Time PCR Detection System. Equivalent amount of MAGwise Paramagnetic Beads was added to the amplified cDNA in each well of the 96 well plate. The cDNA was allowed to bind for 5 minutes. The plate was placed on a magnet to allow the beads to pellet on the inner walls of each well. Supernantant was discarded and the beads were washed once using 80% ethanol. The cDNA was eluted using 20uL of 10mM Tris solution. Purified cDNA was stored at -20^°^C for short term storage.

### Picogreen assay

As per plexWell protocol, we quantified the cDNA concentration of a subset of samples using th eQuant-iT Picogreen dsDNA Assay kit (Thermo Fisher, Waltham, MA, USA). Standards ranging from 25pg/mL to 25ng/mL were prepared using the Lambda Standard DNA (100ug/mL) provided in the kit. 1uL of prepared dsDNA was diluted in 99uL of 1X TE buffer to get a 1:100 dilution. A 200-fold aqueous Quant-iT Picogreen working solution was prepared in 1X TE buffer at a volume of 100uL/well. The standards and samples were plated in a flat bottom Corning 96 well plate using a 125-1250uL multi-channel pipette. Quant-iT Picogreen working solution added at 100uL/well using a 125-1250uL multi-channel pipette. The samples were incubated at room temperature for 5 minutes, protected from light. The fluorescence from Quant-iT picogreen working solution was read on SpectraMax M5e microplate reader (Molecular Devices, San Jose, CA). The preset protocol on SoftMaxPro software version 5.4 for Picogreen Assay for Nucleic acid was used for analysis.

As per the plexWell protocol, 6 samples per 96 well plate were analyzed on Agilent High Sensitivity DNA Bioanalyzer (up to 15000 base pairs) at the University of Michigan Advanced Genomics Core. The analyzed data showed electropherograms with a summary of fragment sizes. The electropherogram for submitted samples were compared to the example electropherograms provided in the plexWell protocol to check for fragment size discrepancies. A pool of 22 random samples was submitted for QuBit analysis at the Advanced Genomics Core per round of experiment to obtain an average concentration. According to the plexWell protocol, the reagents in the kit were formulated to tolerate up to 10-fold difference in sample input across 96 samples. After fragment distribution and concentration checks, we calculated a global dilution factor using the QuBit concentrations (one per library preparation) as described in the plexWell protocol. This global dilution factor gives the volume for dilution of cDNA prior to tagmentation. 4uL of prepared cDNA was diluted in the calculated Tris-HCl volume.

### Library Preparation

Post dilution, 4uL of the diluted cDNA per sample was used for library preparation. Each sample was individually barcoded in a hard skirted Sample Barcode plate provided in the plexWell 384 Rapid Single Cell RNA Library Prep Kit (SeqWell, Beverly, MA, USA). Each sample was labeled with an i7 index, referred to as a Sample Barcode, using a tagmentation reaction. Post i7 tagging, the 96 samples were pooled to a final volume of 800-860uL. Equivalent amount of MAGwise Paramagnetic Beads was added to the pooled SB (Sample Barcode) reactions. The cDNA was allowed to bind to the beads for 5 minutes. The beads then formed a pellet on the inner wall of the tube, as it was placed on a magnetic stand.

The pellet was washed two times with 80% Ethanol. 40uL of 10mM Tris was used to elute out purified SB reaction pool. Next, the purified SB reaction pool was labelled with an i5 index, referred to as a Pool Barcode, using a tagmentation reaction. Post i5 tagging, an equivalent amount of MAGwise Paramengnetic beads was added to the PB (Pool Barcode) reaction. The DNA was allowed to bind to the beads. The beads then formed a pellet on the magnetic stand and was washed two times using 80% ethanol. Purified PB product was amplified using Bio-Rad CFX96 Touch Real-Time PCR Detection System for 12 cycles. Post amplification, the total volume for the reaction was measured and the DNA was diluted to a total volume of 205uL with 10mM Tris. 200uL of the diluted product was transferred to a new 1.5mL LoBind tubes for purification. 5uL of the unpurified product was stored as control. 0.8 equivalents of MAGwise paramagnetic beads were added to the PB product and DNA was allowed to bind to the beads. The beads were allowed to form a pellet on the inner wall of the tube. The pellet was washed two times using 80% ethanol. Purified library was eluted using 32uL of 10mM of Tris. 28uL of the purified product was transferred to a new 1.5mL of LoBind tube. 4 libraries were prepared, each having a unique Pool Barcode. Purified libraries were stored at -20^°^C for short term storage. Library QC was done on the Agilent Bioanalyzer (High Sensitivity DNA 5000 kit) at the Advanced Genomics Core.

### RNA-Sequencing and Data Processing

Libraries were sequenced at Illumina NovaSeq 6000 l at the Advanced Genomics Core. FASTQ reads were demultiplexed back into individual samples based on the i7 and i5 indices. Sequencing data were transferred to the University of Michigan Great Lakes high performance computing cluster for analysis. Sequencing read quality was assessed via FastQC and MultiQC. Reads were aligned to a splice junction aware build of the human genome (GRCh38) using STAR. Aligned reads were assigned to genes using featureCounts, where multimapping and multi-overlapping reads were not counted.

### Differential Gene Expression Testing

Read count matrices from featureCounts were loaded into edgeR, and samples with fewer than 1,000,000 mapped reads were excluded from downstream analysis. Genes with low expression were excluded from analysis using the edgeR filterByExpr() function with default settings. Normalization factors and dispersion were calculated prior to generating a log2-transformed normalized counts per million (cpm) matrix for downstream analysis. Differential gene expression between each treatment and the relevant controls was calculated using quasi-likelihood negative binomial generalized log-linear modeling in edgeR. Genes were considered differentially expressed between a treatment and control at a false discovery rate (FDR) adjusted p-value less than 0.05.

### Benchmark Dose Analyses

Gene specific benchmark doses and best fit benchmark dose model were identified using BMDExpress v.2.3, a free software package developed by the EPA and available for download (https://github.com/auerbachs/BMDExpress-2/releases). Best practices for dose-response modeling for each chemical were conducted according BMDExpress online documentation (https://bmdexpress-2.readthedocs.io/en/feature-readthedocs/basic_workflow/). Normalized counts per million reads were imported into BMDExpress and prefiltered using One-Way ANOVA for significance at a p-value of < 0.05 to identify genes showing significant increasing or decreasing concentration responses. To determine concentration-response relationships, the filtered data were modeled in BMDExpress with hill, power, linear, polynomial (2°, 3°, 4°), and exponential (2, 3, 4, 5) models and the best fit model was chosen. The benchmark response (BMR) was set to 1 standard deviation relative to control response, maximum iterations of the model was set to 250, a confidence level of 0.95 was used, and a power was restricted to greater than or equal to 1 for the power model. Best fit models were chosen for each dose response relationship per gene using nested chi square to select for the best model followed by lowest Akaike information criterion (AIC). Hill models were flagged if its ‘k’ parameter was smaller than the lowest positive dose and flagged hill models were included in the data. Benchmark concentrations with values above the highest noncytotoxic dose (25 µM) were excluded from further analysis. We conducted “Defined Category” analyses to additionally assess the dose-response relationship to the Molecular Signature Database “Hallmark” gene sets, embryonic stem cell genes, breast cancer molecular signatures, and breast stem cell genes.^24–27^ All original expression data, ANOVA filtered results, BMD results, and subsequent BMDExpress analysis for all chemicals are included in a .bm2 file available in supplementary information.

For hypergeometric enrichment testing of BMD data, we used the R package HypeR.^28^ Gene sets were derived from Molecular Signatures Database’s “Hallmark” and “CGP” (Chemical and Genetic Perturbations) gene sets and user-defined gene sets from Pal et al. (2021) and Grashow et al. (2018).^29,30^ Background gene expression included all human genes (23467). Results were filtered for significance at an FDR of < 0.05 and visualized by heatmap using heatmap.2 function from the R package gplots.

### Comparison of NHANES exposure levels to benchmark doses

We compared bioactive doses of the assayed chemicals to human biomarker concentrations measured as part of the National Health and Nutrition Examination Survey following our previously established methodology.^21^ NHANES is designed to understand the health status of the US population, including a substantial number of chemical biomarker measurements in urine, blood, and serum. For this analysis, we used our curated NHANES dataset, which contains information on chemical biomarker concentrations of the 21 assessed chemicals in up to 57786 women recruited from 1999-2018.^31^ Linkage between chemicals used *in vitro* and their corresponding exposure biomarkers in NHANES are shown in Supplemental Table 2. Chemical biomarker concentrations were converted to molarity units and then compared to the benchmark dose concentrations identified through the benchmark dose analyses of the RNA-seq data using boxplots. These comparisons are designed to help us understand whether dysregulated gene expression due to toxicant exposure may be occurring at population relevant levels. Overlapping NHANES biomarker concentrations and BMD concentrations indicate that chemical biomarkers measured in US women in NHANES occur within the estimated benchmark doses.

### Secondary analysis of normal breast single cell data for cell type composition prediction

To quantify whether treatments altered the cellular composition of our samples, we performed bioinformatic deconvolution of our bulk RNA-seq data based on a single cell atlas of the normal human mammary gland (Pal et al., 2021), where 10 normal mammary gland samples were processed for single cell analysis using the 10x Genomics Chromium platform.^29^ Sample specific count matrices of the single cell RNA-seq data were downloaded from the Gene Expression Omnibus and processed via Seurat. Briefly, we excluded cells which express fewer than 200 or more than 5000 genes, along with cells with greater than 15% mitochondrial reads. Data were log-normalized and scaled, and then clustered with 0.02 resolution, following Pal et al’s methods, to identify four cell clusters. Clusters were then assigned to “luminal progenitor”, “mature luminal”, and “myoepithelial” identities based on marker gene expression. Gene expression signatures for each of the cell types were calculated using the FindAllMarkers() function in Seurat.

We then used these single cell gene expression data to estimate cell type proportions in our data using the Multi-subject Single Cell (MuSiC) deconvolution method. MuSiC predicts cell type proportions in bulk RNA-seq data based on annotated single cell RNA expression data from multiple donors. Using MuSiC, we estimated the proportions of myoepithelial, mature luminal, and luminal progenitor cells in each sample to test the hypothesis that chemical treatments may induce cellular plasticity or differences in cell type marker expression. Differences in cell type proportions by chemical dose were calculated with generalized linear modeling in R, comparing back to the relevant control samples (DMSO or water) as the reference group. Differences in estimated cell type proportions by treatment were compared to control using generalized linear modeling.

### Data Availability

The sequencing data have been deposited at the Gene Expression Omnibus (accession GSE220051; reviewer token shyjiiikjnabbeh).

## Results

To understand the impacts of exposure disparities chemicals on dysregulated breast cancer biology, we treated non-tumorigenic MCF10A cells with chemicals found through previous analysis of human population biomonitoring data that represent exposures from metals, pesticides, personal care products, and other industrial and environmental sources, including per- and polyfluoroalkyl substances.^19,21^ After filtering expression data for genes with significant alterations, fold change and significance of gene expression varied substantially by chemical and by dose (Figure 1, Table 1, Supplemental Figure 1, Supplemental Table 3). For example, p,p’-DDE showed over 700 differentially expressed genes (DEGs) at all 4 doses, while mercury only had over 10 DEGs at the highest dose of 25 µM. Arsenic and cadmium both had DEGs at all four doses, but had over 8000 and 4000 DEGs at 25 µM, respectively. As an example of the differences in effect of each chemical at a given dose, volcano plots showing the results of differential expression analysis comparing the 2.5 µM dose to control are shown in Figure 1, with the comparisons for the other doses compared to control included in Supplemental Figure 1.

**Table 1:** Differentially expressed genes (DEGs) (FDR < 0.05 relative to control) by dose for each chemical.

|  | 25nM vs. Ctrl |  | 250nM vs. Ctrl |  | 2.5uM vs. ctrl |  | 25uM vs. Ctrl |  |  |
| --- | --- | --- | --- | --- | --- | --- | --- | --- | --- |
| Chemical | Up | Down | Up | Down | Up | Down | Up | Down | Total DEGs |
| Arsenic | 8 | 13 | 41 | 34 | 137 | 211 | 3604 | 4701 | 8749 |
| Mercury | 0 | 0 | 0 | 1 | 0 | 4 | 6 | 10 | 21 |
| Copper | 9 | 17 | 72 | 53 | 246 | 182 | 288 | 467 | 1334 |
| Cadmium | 1 | 4 | 0 | 7 | 0 | 67 | 2539 | 2087 | 4705 |
| Lead | 0 | 15 | 29 | 48 | 78 | 85 | 235 | 343 | 833 |
| MPB | 8 | 12 | 30 | 39 | 32 | 24 | 111 | 60 | 316 |
| PPB | 5 | 21 | 5 | 23 | 3 | 10 | 1 | 0 | 68 |
| BPA | 52 | 13 | 20 | 20 | 98 | 16 | 12 | 1 | 232 |
| BPS | 20 | 16 | 3 | 10 | 2 | 3 | 5 | 4 | 63 |
| BPF | 6 | 15 | 12 | 9 | 0 | 22 | 5 | 2 | 71 |
| Phenanthrene | 59 | 5 | 72 | 6 | 12 | 4 | 7 | 3 | 168 |
| PFOA | 96 | 78 | 92 | 53 | 56 | 66 | 52 | 72 | 565 |
| PFDA | 65 | 5 | 32 | 22 | 38 | 2 | 57 | 6 | 227 |
| PFNA | 20 | 2 | 24 | 4 | 76 | 3 | 68 | 3 | 200 |
| PFOS | 21 | 6 | 17 | 3 | 21 | 7 | 41 | 5 | 121 |
| Thiram | 0 | 1 | 3 | 5 | 0 | 32 | 2521 | 2549 | 5111 |
| DCP_25 | 0 | 0 | 0 | 0 | 0 | 1 | 0 | 2 | 3 |
| DCB_14 | 0 | 0 | 0 | 0 | 1 | 0 | 1 | 0 | 2 |
| PCB153 | 1 | 0 | 1 | 50 | 0 | 0 | 1 | 0 | 53 |
| PCB187 | 1 | 1 | 0 | 0 | 0 | 0 | 4 | 1 | 7 |
| p,p'-DDE | 467 | 312 | 852 | 572 | 645 | 562 | 960 | 749 | 5119 |
|  |  |  |  |  |  |  |  |  | 27968 |

**Figure 1:**
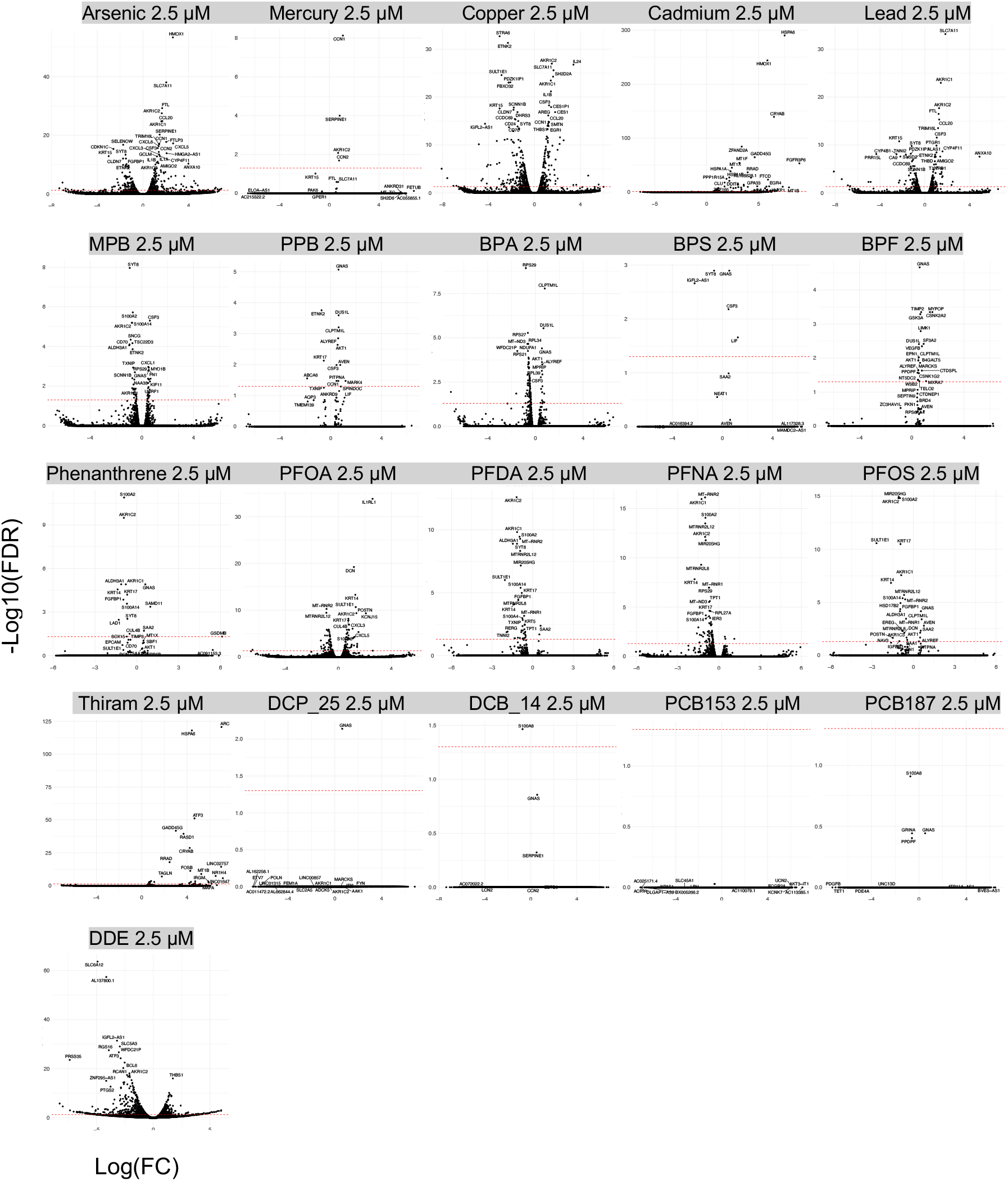
Differential gene expression for 21 disparities associated chemicals at 2.5 µM. Differential gene expression between each chemical dose and controls was calculated using quasi-likelihood negative binomial generalized log-linear modeling in edgeR. Red lines mark a false discovery rate (FDR) adjusted p-value of 0.05.

Genes related to breast cancer were commonly found to be differentially expressed by this panel of chemicals at various doses. Of note, G protein alpha subunit (GNAS), a gene related to breast cancer cell proliferation and migration, expression changes were observed across 28 exposures.^32^ Additionally, aldo-keto reductases AKR1C1, AKR1C2, and AKR1C3 expression changed across 30, 38, and 6 exposures, respectively, and these genes are associated with hormone signaling and breast cancer proliferation.^33^ Keratins KRT14, KRT15, and KRT17 expression levels also changed across 17, 9, and 18 exposures, and their dysregulation is associated with breast and other cancers.^34-36^ Serine protease inhibitor, clade E member 1 (SERPINE1), a gene associated with TNBC chemotherapy resistance, also showed changes in expression across 12 chemical treatments.^37^

To characterize the dose-response relationship of each exposure on gene expression, we performed benchmark dose modeling, to define the range of benchmark doses for the different chemicals (Figure 2). Best benchmark doses were estimated from a set of dose-response models for each gene for each chemical (Figure 2A). The total number of genes filtered by ANOVA and fit to benchmark models was different for each chemical, with some chemicals (arsenic, cadmium, and thiram) inducing dose-responsive alterations of over 1000 different genes.

**Figure 2:**
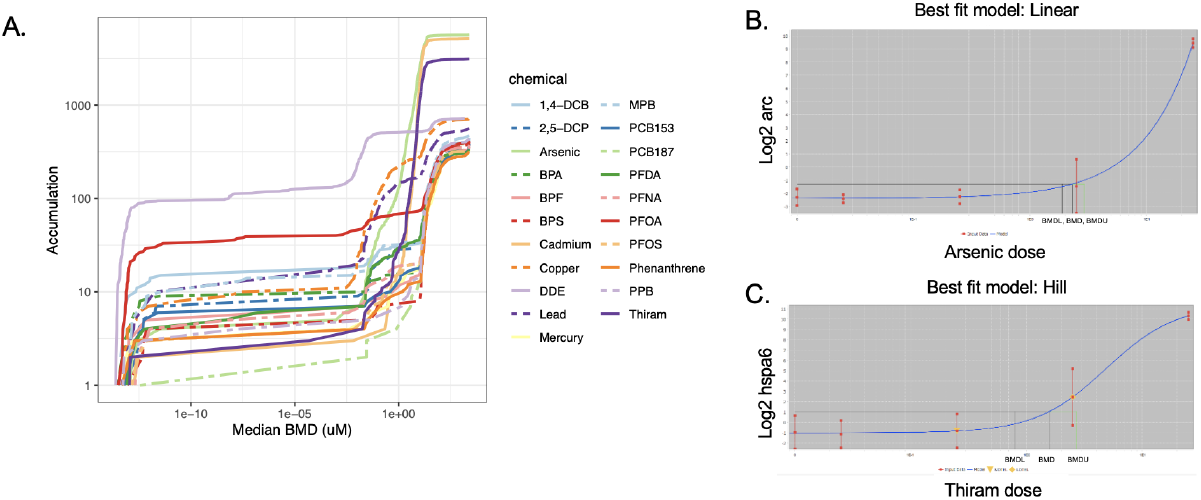
Benchmark dose accumulation plots and individual benchmark dose modeling graphs. In the accumulation plot of median best benchmark doses (A), each plotted point represents a differentially expressed gene and each differently colored line represents individual chemicals. On the right, example individual best fit benchmark dose graphs are shown for the gene *ARC* in arsenic treated cells (B) and for the gene *HSPA6* in thiram treated cells (C). For B and C, experimental data is shown in red, best fit model in blue, and lower bound of the 95% confidence interval of the benchmark dose (BMDL), benchmark dose (BMD), and upper bound of the 95% confidence interval of the benchmark dose (BMDU) labeled on each graph.

Dose-response fit was assessed via examination of the best fit dose-response curves for individual gene/chemical combinations. Two example gene BMD curves are displayed in Figure 2 B and C, showing the gene expression of *ARC* following exposure to a range of arsenic doses in a linear best fit model (Figure 2B) and thiram’s dose-response regulation of gene *HSPA6* in a hill best fit model (Figure 2C) across the range of treated dose. All benchmark dose response data for each dysregulated gene for every chemical tested is included in the supplementary bm2 file.

We next wanted to contextualize the doses required *in vitro* to dysregulate gene expression with biomarker levels detectable women in the United States. We compared benchmark dose results from our *in vitro* experiments to National Health and Nutrition Examination Survey (NHANES) biomarker concentration data for each chemical, plotting the range of blood or urine chemical biomarker concentrations in US women compared to the range of BMDs for each chemical from our *in vitro* data (Figure 3). For many of the chemicals, the levels detectable in the US population are sufficient to alter gene expression in MCF10A cells, reflected by an overlap in the distributions of NHANES biomarker concentrations and benchmark doses for a given chemical. For many chemicals (phenanthrene, PFNA, PFDA, PCB187, mercury, cadmium, bisphenol S, bisphenol F, bisphenol A, arsenic), chemical biomarker concentration in the population were below the vast majority of benchmark doses for altered gene expression. Conversely, for a number of chemicals, population biomarker concentrations were comparable to, or above, the bulk of the benchmark doses. These chemicals included propyl paraben, PFOA, p,p’-DDE, methyl paraben, lead, copper, and 2,5-dichlorophenol.

**Figure 3:**
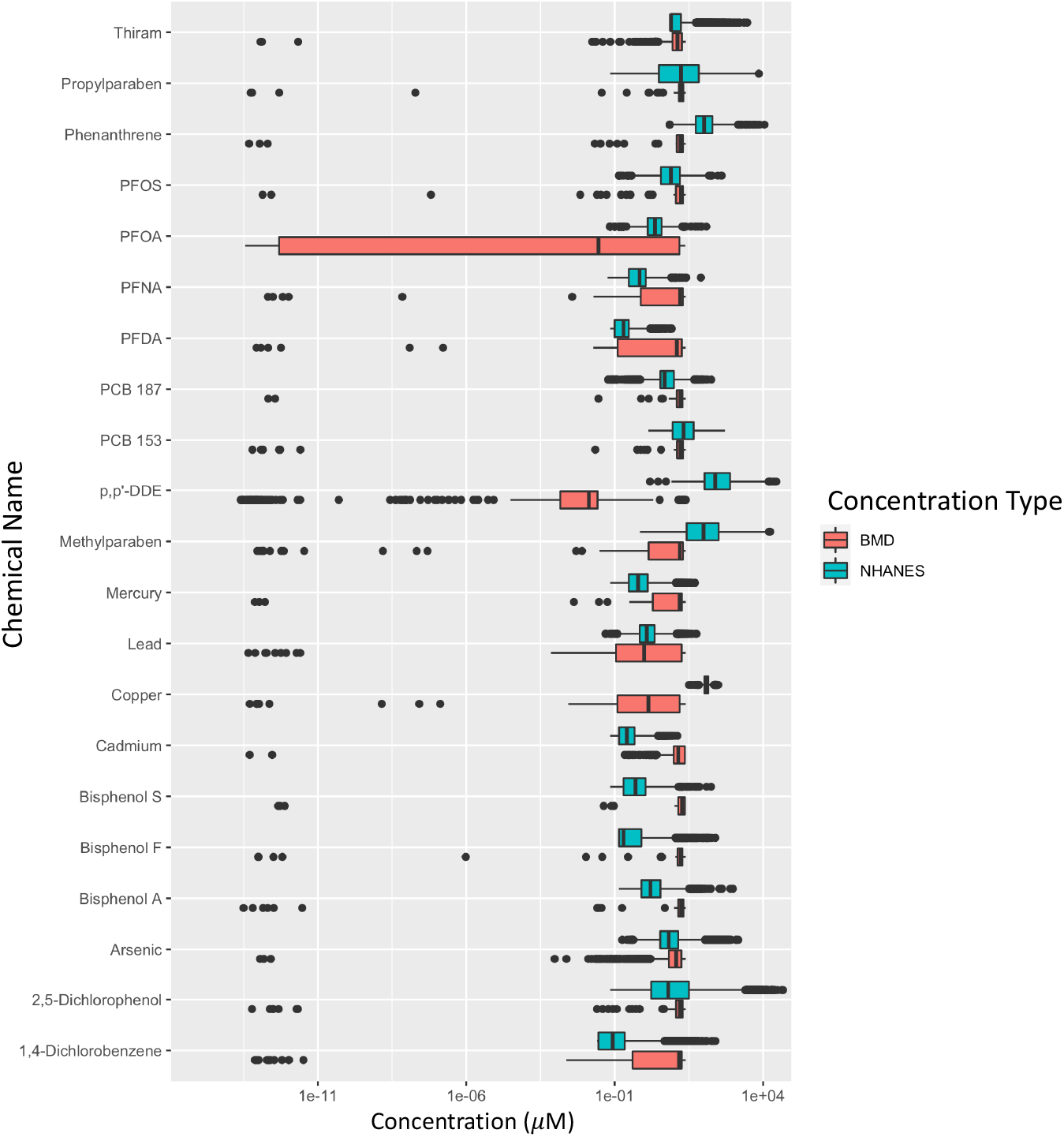
Comparison of benchmark dose modeling results with NHANES exposure biomarker levels. Median BMD for each differentially expressed gene (red) and NHANES chemical biomarker concentration data, converted to molarity units, for female participants (blue) plotted for each chemical.

We next wanted to quantify the biological processes altered by chemical exposure and characterize the doses at which these processes are dysregulated. BMD modeled genes were assessed using MSigDB’s Hallmark gene sets consisting of 50 biological processes. The supplemental bm2 file contains benchmark dose gene data for every hallmark category for each chemical organized under “functional classifications.” Figure 4A displays an accumulation plot depicting the median BMD at which Hallmark gene sets were significantly impacted. The accumulation plot demonstrates low dose effects present for many of the Hallmarks. Figure 4B shows a heatmap of significantly changed hallmark processes - 9 of the chemicals showed significant enrichment or downregulation for 1 or more of the Hallmark gene sets. Notably, many of these processes are changes that are related to carcinogenesis, including the epithelial mesenchymal transition, inflammatory response, and reactive oxygen species pathway. Epithelial to mesenchymal transition was upregulated by arsenic, cadmium, copper, lead, and thiram at median BMD dose of 7.58 µM, 9.83 µM, 0.60 µM, 0.55 µM, and 9.01 µM, respectively.

**Figure 4:**
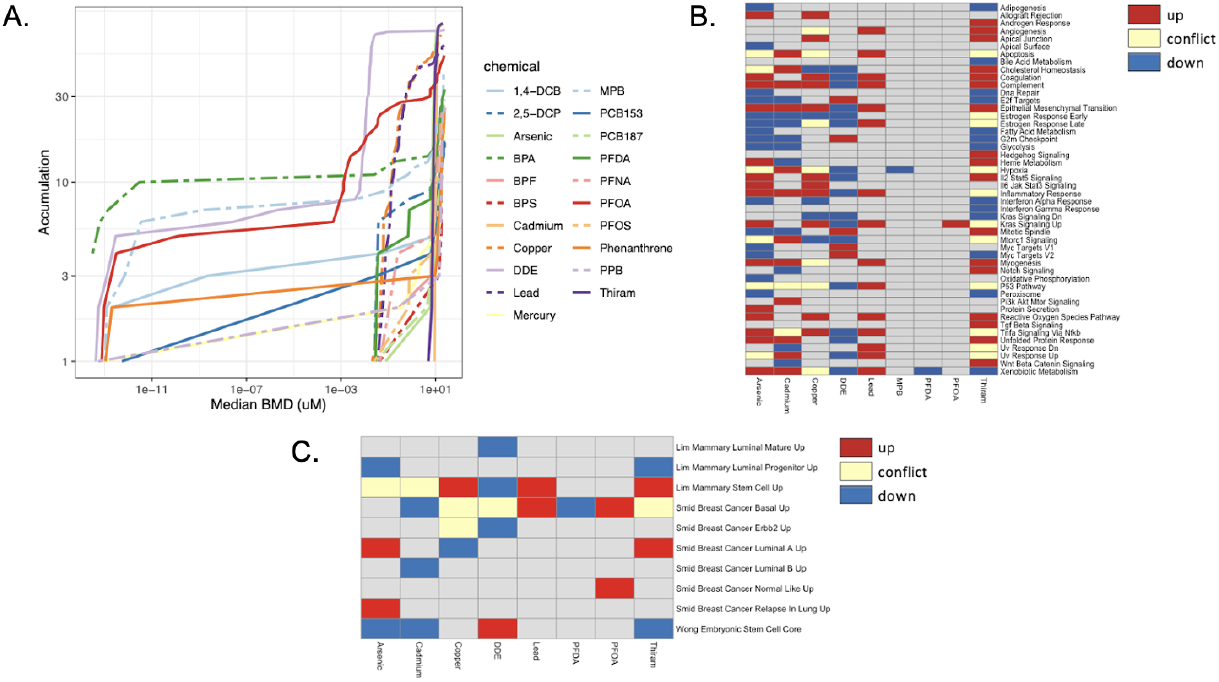
Enrichment analyses. All of MSigDB’s “Hallmark” biological processes as regulated by each chemical across different median BMDs (A). Nine of the chemicals showed significant enrichment or downregulation for 1 or more of 50 biological processes defined by hallmark gene sets (B). Eight chemicals showed significant alterations of breast cancer or stemness specific gene pathways as defined by some of MSigDB’s chemical and genetic perturbations sets (C).

Inflammatory response was upregulated by arsenic, cadmium, copper, and lead at median BMD dose of 8.26 µM, 12.28 µM, 0.27 µM, and 0.19 µM, respectively. Reactive oxygen species was upregulated by arsenic, copper, lead, and thiram at median BMD dose of 8.31 µM, 0.18 µM, 0.12 µM, and 10.59 µM, respectively. p,p’-DDE downregulated epithelial mesenchymal transition genes (median BMD of 0.0079 µM) and upregulated cell cycle related processes, including E2f targets and G2m checkpoint (median BMD of 0.011 µM and 0.015 µM, respectively), both of which were downregulated by arsenic, cadmium, and thiram (median BMDs of 14.09 µM, 10.77 µM, and 8.27 µM, respectively, for E2f targets and of 8.91 µM, 10.68 µM, and 8.96 µM, respectively, for G2m checkpoint), highlighting the potential for distinct mechanisms of promoting carcinogenesis depending on the chemical involved.

To assess specific pathways related to breast cancer, additional enrichment analyses assessed additional breast cancer associated biological pathways in MSigDB’s Chemical and Genetic Perturbations (CGP) gene sets (Figure 4C and Supplemental Figure 1): Gene sets from three studies related to breast cancer and stemness.^25-27^ Eight chemicals differentially regulated these features at various doses. Genes upregulated in mammary stem cells were found upregulated by copper, lead, and thiram (median BMDs of 0.59 µM, 0.40 µM and 11.78 µM, respectively) and genes upregulated in embryonic stem cells were upregulated by p,p’-DDE (median BMD of 0.012 µM). Additionally, genes upregulated in basal subtypes of breast cancer were found upregulated by lead and PFOA (median BMDs of 0.49 µM and 0.0026 µM), while genes upregulated in luminal A breast cancers were found upregulated by arsenic and thiram (median BMD of 10.36 µM and 8.98 µM). PFOA also upregulated genes that are upregulated in normal like breast cancers (median BMD of 0.00093 µM) and arsenic upregulated genes upregulated in breast cancer relapse in the lung (median BMD of 10.16 µM).

To assess potential breast cancer specific alterations regulated by our chemicals of interest, we tested for enrichment with a publicly available breast carcinogenesis gene panel, BCScreen.^30^ The overlapping genes within different categories of breast carcinogenesis are shown in supplemental table 4. The gene panel includes 500 genes total divided into 14 categories, with 33 genes per category other than the “mammary” category which includes 71 genes. Of these, arsenic, cadmium, and thiram showed BMD gene alterations in all 14 categories, and copper, lead, and p,p’-DDE showed at least one gene alteration in most categories.

We next wanted to test the hypothesis that disparities associated chemicals induce changes consistent with a luminal-to-basal transition, a process likely important in the development of basal breast cancers. We performed cell type deconvolution based on a normal breast single cell RNA-seq reference to predict the proportions of myoepithelial, mature luminal, and luminal progenitor cells in each of our exposures (Figure 5 and Supplemental Figures 2-4). ^29^ Interestingly, PFOA showed a significant decrease in myoepithelial cell proportion from around 65% to 52% across 3 doses (Supplemental Figure 2). Arsenic, copper, and lead showed a significant increase in the proportion of myoepithelial cells across all doses compared to control, while p,p’-DDE showed a significant increase in this proportion at the 3 highest doses (Figure 5). These data suggest that these chemicals induce an expression signature consistent with a luminal-to-basal transition, a characteristic of aggressive basal-like breast cancers.

**Figure 5:**
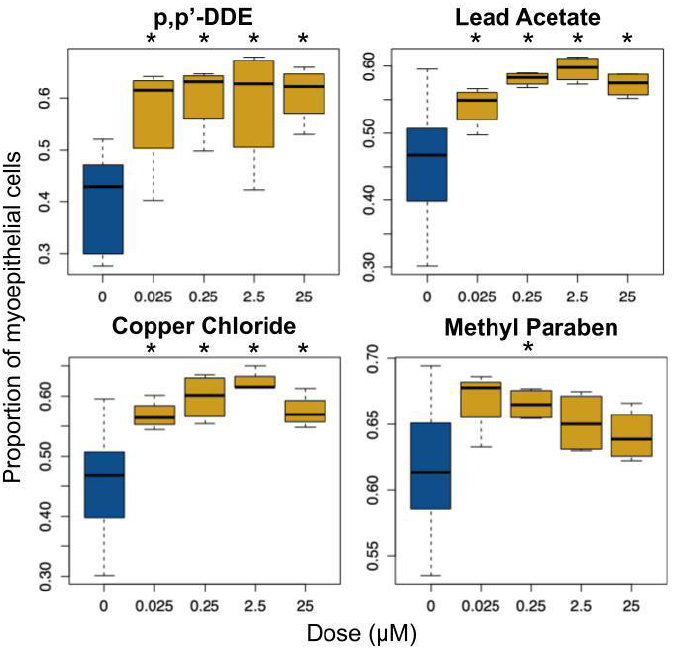
MuSiC deconvolution results. Comparing the proportion of estimated myoepithelial cells in MCF10A cultures treated with varying doses of arsenic (A), copper (B), lead (C), and p,p’-DDE (D) all showed significant increases in myoepithelial cell proportions relative to control.

## Discussion

Across all 21 chemicals representing exposures commonly affecting women in the United States, nearly 30,000 differentially expressed genes were found dysregulated in exposed breast cells compared to controls. While many of these alterations were found at the highest doses, lower dose effects were also seen, with median BMDs falling near the range of NHANES biomarker levels for all chemicals. This suggests that the gene expression effects seen in this study are representative of effects seen in humans, and so our analysis may link chemical exposures to incidence and characteristics of breast cancer. Interestingly, of the chosen set of chemicals, only nine have been classified by IARC as possible, probable, or definite carcinogens to humans, making this study both novel and relevant in assessing underlying chemical influence on the biology of cancers.

Specific gene alterations included some commonalities across chemical treatments showing expression changes in genes related to breast cancer progression. This included GNAS, aldo-keto reductases, keratins, and SERPINE1. GNAS has been implicated in breast cancer cell proliferation and epithelial to mesenchymal transition via modulation of the PI3K/AKT/Snail1/E-cadherin signaling axis.^32^ Aldo-keto reductases are involved in estrogen and progesterone production and are thus involved in pro-proliferative signaling of hormone responsive tumors.^33^ Dysregulation of keratin expression levels has also been associated with breast cancer, with increased KRT14 expression associated with invasiveness, decreased KRT15 expression associated with poor prognosis, and increased KRT17 associated with the development of TNBC.^34-36^ SERPINE1 is associated with resistance of TNBC to paclitaxel and knockdown of SERPINE1 reversed resistance of breast cancer to paclitaxel through downregulation of vascular endothelial growth factor A (VEGFA). ^37^

In addition to common gene expression changes across chemical treatments, our enrichment analyses suggest that chemicals alter breast cell biology by several distinct mechanisms. Metals including arsenic, cadmium, copper, and lead were found to induce epithelial to mesenchymal transition, inflammatory response, and reactive oxygen species pathways, consistent with previous literature on metal-induced carcinogenesis that places particular emphasis on oxidative stress related carcinogenesis.^38–40^ In breast cancer, arsenic and cadmium have been associated with oxidative stress related pathways of malignant transformation.^41^ While these metals as well as many of the other chemicals in our analysis are known to play an endocrine disrupting role that could lead to breast tumorigenesis, these findings propose alternative routes of carcinogenesis.^42–44^ p,p’-DDE downregulated epithelial to mesenchymal transition in our study but altered cell cycle processes. Previous literature has found that a mixture of organochlorine pesticides including p,p’-DDE had potential to modulate proliferation of breast cancer cell lines partially through induction of cell cycle entry.^45^

Alterations in stemness features were also seen across eight chemicals analyzed, important because the cancer stem cell hypothesis states that there is a small subset of cancer stem cells that result in cancer initiation and invasion, and so alterations in stemness features can lead to increased tumorigenicity and metastasis.^46^ We found stemness signatures enhanced by copper, lead, thiram, and p,p’-DDE, findings that could indicate novel mechanisms of chemically induced carcinogenesis.

These findings may be particularly relevant in the context of breast cancer disparities. We previously identified that in the US, non-Hispanic Black women are disproportionately exposed to many of the chemicals assayed here.^19^ Non-Hispanic black women are also three times more likely to be diagnosed with triple negative breast cancers and approximately 40% more likely to die from a breast cancer diagnosed compared to white women.^47,48^ Triple negative breast cancers often have a basal-like phenotype and may derive from a luminal to basal cell state transition.^14-16^

Here using enrichment analysis, we found upregulation of basal subtype genes by lead and PFOA and cell type proportion analysis found upregulated basal proportions by arsenic, copper, lead, and p,p’-DDE and a decrease in basal cell proportions by PFOA. Previous literature has shown that arsenic is capable of inducting transition from luminal to basal features, but this association has not been studied for the other chemicals.^49^ These findings highlight an increased need to examine chemical exposures as a modifiable risk factor for aggressive breast cancers, particularly in the context of breast cancer disparities.

One major limitation of the present study is that all exposures were done on an acute timescale (48 hours) while actual human exposures to the chemicals analyzed here are likely chronic, over the course of years or a lifetime. Another limitation is that only one cell type was used for this study, normal breast cells taken from a white woman. Ongoing work is considering treatment on a more chronic timescale as well as different breast cell lines representing diverse donors.

Additionally, future work could consider treatment of diverse breast cancer cell lines, ranging from more basal to more luminal cell types to investigate changes in luminal or basal features related to chemical exposure. Another limitation in our analyses integrating benchmark dose levels with NHANES biomarker concentrations is that the NHANES chemical biomarkers are measured in urine and blood. These levels may or may not reflect mammary tissue levels. As additional research is conducted on biomonitoring of mammary tissues, or physiologically based toxicokinetic models reflecting mammary gland concentrations are developed, these data and methods will substantially inform our understanding of how chemical exposures impact breast cancer risk.

Overall, our transcriptomic analysis uncovers molecular mechanisms by which chemicals induce more aggressive forms of breast cancer. This includes both features related to known carcinogenic properties of these chemicals as well as potentially novel mechanisms, opening the door to future avenues of investigation to elucidate how different exposures relate to breast carcinogenesis and breast cancer disparities.

## Acknowledgements

This work was supported by grants from the National Institutes of Health (R01 ES028802, R01 AG072396, T32 ES 007062, P30 ES017885, P30 CA046592).

