## Supplemental Tables and Figures for "High Throughput Transcriptomics to Understand Chemical Drivers of Aggressive Breast Cancer Subtypes"

| Chemical | CAS number | Vendor; Catalog number | Solvent |
|---|---|---|---|
| Lead (II) Acetate Trihydrate | 6080-56-4 | Sigma Aldrich; 316512 | Water |
| Copper (II) Chloride | 7447-39-4 | Sigma Aldrich; 222011 | Water |
| Sodium (meta) Arsenite | 7784-46-5 | Sigma Aldrich; S7400 | Water |
| Mercury (II) Chloride | 7487-94-7 | Sigma Aldrich; 215465 | Water |
| Cadmium Chloride | 10108-64-2 | Sigma Aldrich; 202908 | Water |
| Bisphenol A | 9980-05-7 | Chem Service; S-F7295S | Dimethyl sulfoxide (DMSO) |
| Bisphenol S | 80-09-1 | Sigma Aldrich; 103039 | Dimethyl sulfoxide (DMSO) |
| Bisphenol F | 620-92-8 | Sigma Aldrich; 51453 | Dimethyl sulfoxide (DMSO) |
| Methylparaben | 99-76-3 | Sigma Aldrich; 47889 | Dimethyl sulfoxide (DMSO) |
| Propylparaben | 94-13-3 | Sigma Aldrich; 1577008 | Dimethyl sulfoxide (DMSO) |
| Phenanthrene | 85-01-8 | Sigma Aldrich; P11409 | Dimethyl sulfoxide (DMSO) |
| PFOA | 335-67-1 | Sigma Aldrich; 77262 | Water |
| PFDA | 335-76-2 | Sigma Aldrich; 177741 | Dimethyl sulfoxide (DMSO) |
| PFNA | 375-95-1 | Sigma Aldrich; 394459 | Dimethyl sulfoxide (DMSO) |
| PFOS | 1763-23-1 | Sigma Aldrich; 77282 | Dimethyl sulfoxide (DMSO) |
| Thiram | 137-26-8 | Sigma Aldrich; 45689 | Dimethyl sulfoxide (DMSO) |
| 2,5-Dichlorophenol | 583-78-8 | Sigma Aldrich; D70007 | Dimethyl sulfoxide (DMSO) |
| 1,4-Dichlorobenzene | 99106-46-7 | Sigma Aldrich; D56829 | Dimethyl sulfoxide (DMSO) |
| PCB153 | 35065-27-1 | Chem Service; BZ-153 | Dimethyl sulfoxide (DMSO) |
| PCB 187 | 52663-68-0 | Chem Service; BZ-187 | Dimethyl sulfoxide (DMSO) |
| DDE | 72-55-9 | Chem Service; N-10875 | Dimethyl sulfoxide (DMSO) |

**Supplemental Table 1: Chemical information.** 21 chemicals chosen based on NHANES disparity data for this study and their associated CAS number, vendor, catalog number, and solvent in which each was dissolved for cell treatments.

| Chemical Name for In Vitro Treatment | Corresponding NHANES Biomarker | NHANES Chemical Codename | Biomarker Matrix |
|---|---|---|---|
| Lead (II) Acetate Trihydrate | Blood lead (ug/dL) | LBXBPB | Whole Blood |
| Copper (II) Chloride | Serum Copper (ug/dL) | LBXSCU | Serum |
| Sodium (meta) Arsenite | Urinary arsenic, total (ug/L) | URXUAS | Urine |
| Mercury (II) Chloride | Blood mercury, total (ug/L) | LBXTHG | Whole Blood |
| Cadmium Chloride | Blood cadmium (ug/L) | LBXBCD | Whole Blood |
| Bisphenol A | Urinary Bisphenol A (ng/mL) | URXBPH | Urine |
| Bisphenol S | Urinary Bisphenol S (ug/L) | URXBPS | Urine |
| Bisphenol F | Urinary Bisphenol F (ug/L) | URXBPF | Urine |
| Methylparaben | Methyl paraben (ng/ml) | URXMPB | Urine |
| Propylparaben | Propyl paraben (µg/L) | URXPPB | Urine |
| Phenanthrene | 3-hydroxyphenanthrene (ng/L) | URXP05 | Urine |
| PFOA | Perfluorooctanoic acid | LBXPFOA | Serum |
| PFDA | Perfluorodecanoic acid | LBXPFDE | Serum |
| PFNA | Perfluorononanoic acid | LBXPFNA | Serum |
| PFOS | Perfluorooctane sulfonic acid | LBXPFOS | Serum |
| Thiram | Urinary 2-Thioxothiazolidine-4-carboxylic acid (ng/mL) | URXTTC | Urine |
| 2,5-Dichlorophenol | Urinary 2,5-dichlorophenol (µg/L) | URX14D | Urine |
| 1,4-Dichlorobenzene | Blood 1,4-Dichlorobenzene (ng/mL) | LBXVDB | Whole Blood |
| PCB 153 | PCB153 Lipid Adjusted (ng/g) | LBX153LA | Blood (lipid adjusted) |
| PCB 187 | PCB187 Lipid Adjusted (ng/g) | LBX187LA | Blood (lipid adjusted) |
| p',p'-DDE | ppDDE Lipid Adjusted (ng/g) | LBXPDELA | Blood (lipid adjusted) |

**Supplemental Table 2: NHANES Biomarker Information.** Chemicals used *in vitro* and their corresponding exposure biomarkers in NHANES.

***Supplemental Table 3 included as separate excel document***

| | Arsenic | Cadmium | Copper | Lead | Mercury | PFOS | PFOA | PFNA | PFDA | PCB153 | PCB187 | DDE | Thiram | DCB_14 | DCP_25 | MPB | Total by Category |
|---|---|---|---|---|---|---|---|---|---|---|---|---|---|---|---|---|---|
| **Angiogenesis** | 12 | 10 | 6 | 2 | | 1 | 2 | 1 | 1 | 1 | | 2 | 8 | | | | 46 |
| **Apoptosis** | 12 | 10 | 1 | 1 | | | 1 | | | | | 2 | 10 | | | | 37 |
| **Cell Cycle** | 14 | 9 | | | | | 1 | | 1 | | | 4 | 9 | | | | 38 |
| **Epigenetics** | 8 | 10 | 1 | 1 | | | | | | | | 2 | 7 | | | | 29 |
| **Genotoxicity** | 7 | 12 | | | | | | | | | | 3 | | | | | 22 |
| **Growth hormones** | 5 | 5 | | | | | 1 | 1 | | | | | 5 | | | | 17 |
| **Immortalization** | 7 | 9 | 1 | 1 | | | | | | | | 2 | 5 | | | | 25 |
| **Immunomodulation** | 8 | 7 | | | | | | | | | | 2 | 4 | | | | 21 |
| **Inflammation** | 14 | 8 | 4 | 3 | | | | | | | | | 6 | | | | 35 |
| **Mammary** | 25 | 23 | 4 | 3 | | 1 | 1 | | | 1 | 1 | 4 | 11 | 2 | 3 | 1 | 80 |
| **Oxidative stress** | 10 | 6 | 2 | 3 | | | | | | | | | 8 | | | | 29 |
| **Proliferation** | 8 | 11 | 2 | | | | | | | | | 1 | 6 | | | | 28 |
| **Steroid hormones** | 9 | 4 | 5 | 3 | 2 | 1 | 1 | 1 | 2 | | | 3 | 5 | | | | 36 |
| **Xenobiotic metabolism** | 7 | 3 | 2 | 2 | | | | | | 1 | | 3 | 4 | | | | 22 |
| **Total BCScreen** | 146 | 127 | 28 | 19 | 2 | 3 | 7 | 3 | 4 | 3 | 1 | 28 | 88 | 2 | 3 | 1 | 465 |

**Supplemental Table 4: BCScreen enrichment.** Gene alterations overlapping with 14 breast cancer related gene sets as defined by a publicly available breast carcinogenesis gene panel called BCScreen (Grashow et al., 2018; raw gene numbers shown in table).

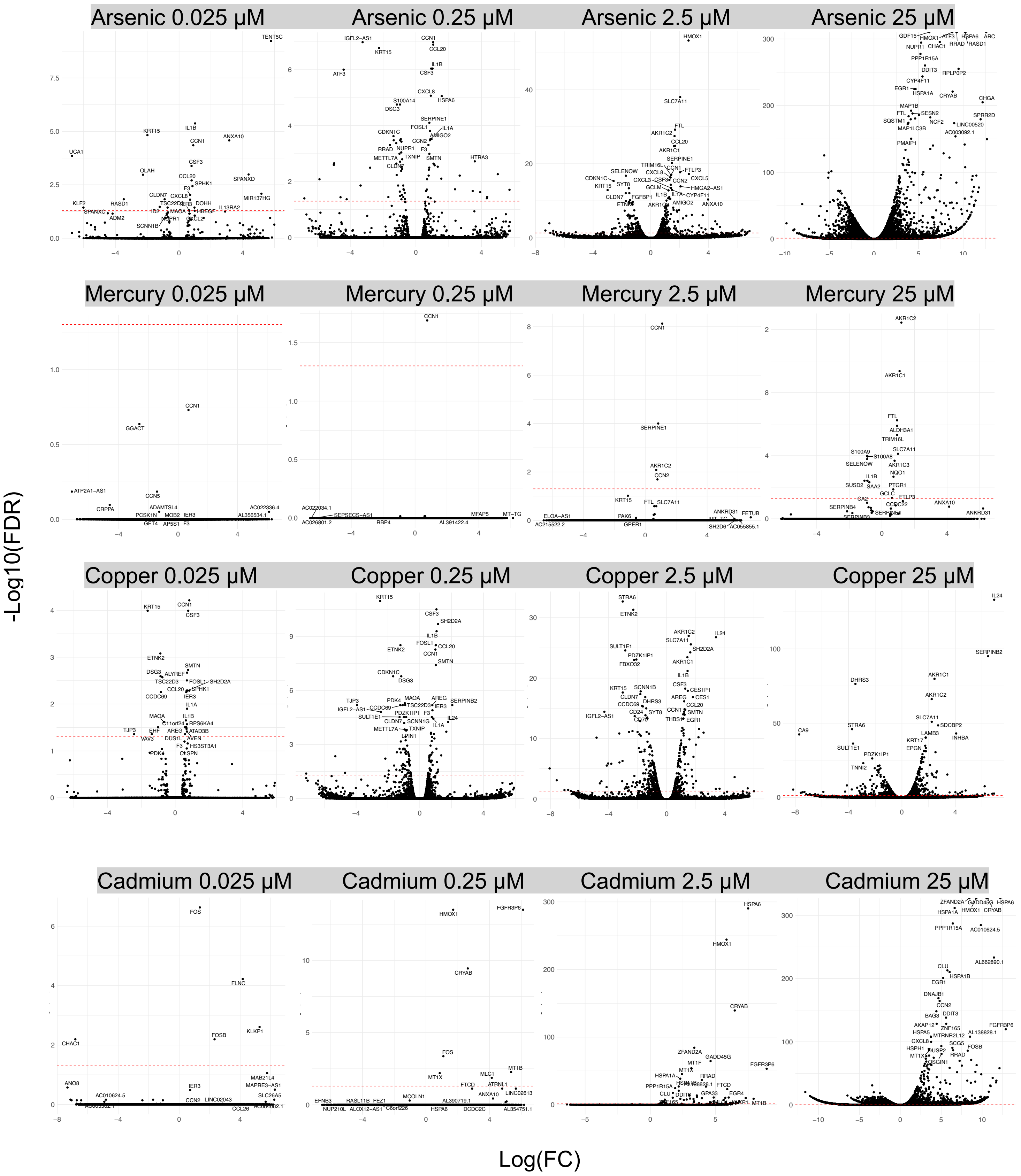
Arsenic 0.025 µM
Arsenic 0.25 µM
Arsenic 2.5 µM
Arsenic 25 µM
Mercury 0.025 µM
Mercury 0.25 µM
Mercury 2.5 µM
Mercury 25 µM
Copper 0.025 µM
Copper 0.25 µM
Copper 2.5 µM
Copper 25 µM
Cadmium 0.025 µM
Cadmium 0.25 µM
Cadmium 2.5 µM
Cadmium 25 µM
-Log10(FDR)
Log(FC)

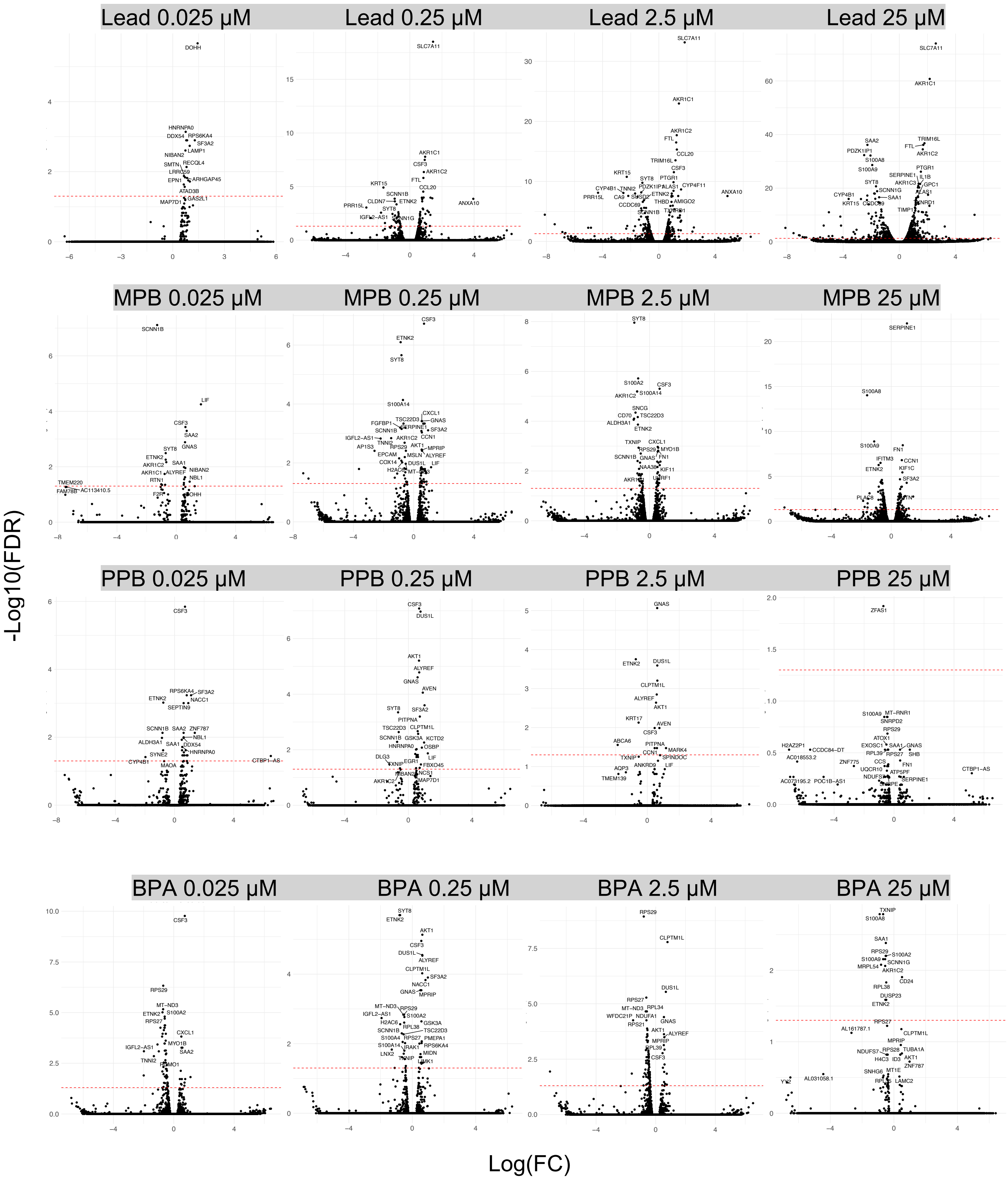
Lead 0.025 μM
Lead 0.25 μM
Lead 2.5 μM
Lead 25 μM
MPB 0.025 μM
MPB 0.25 μM
MPB 2.5 μM
MPB 25 μM
PPB 0.025 μM
PPB 0.25 μM
PPB 2.5 μM
PPB 25 μM
BPA 0.025 μM
BPA 0.25 μM
BPA 2.5 μM
BPA 25 μM
-Log10(FDR)
Log(FC)

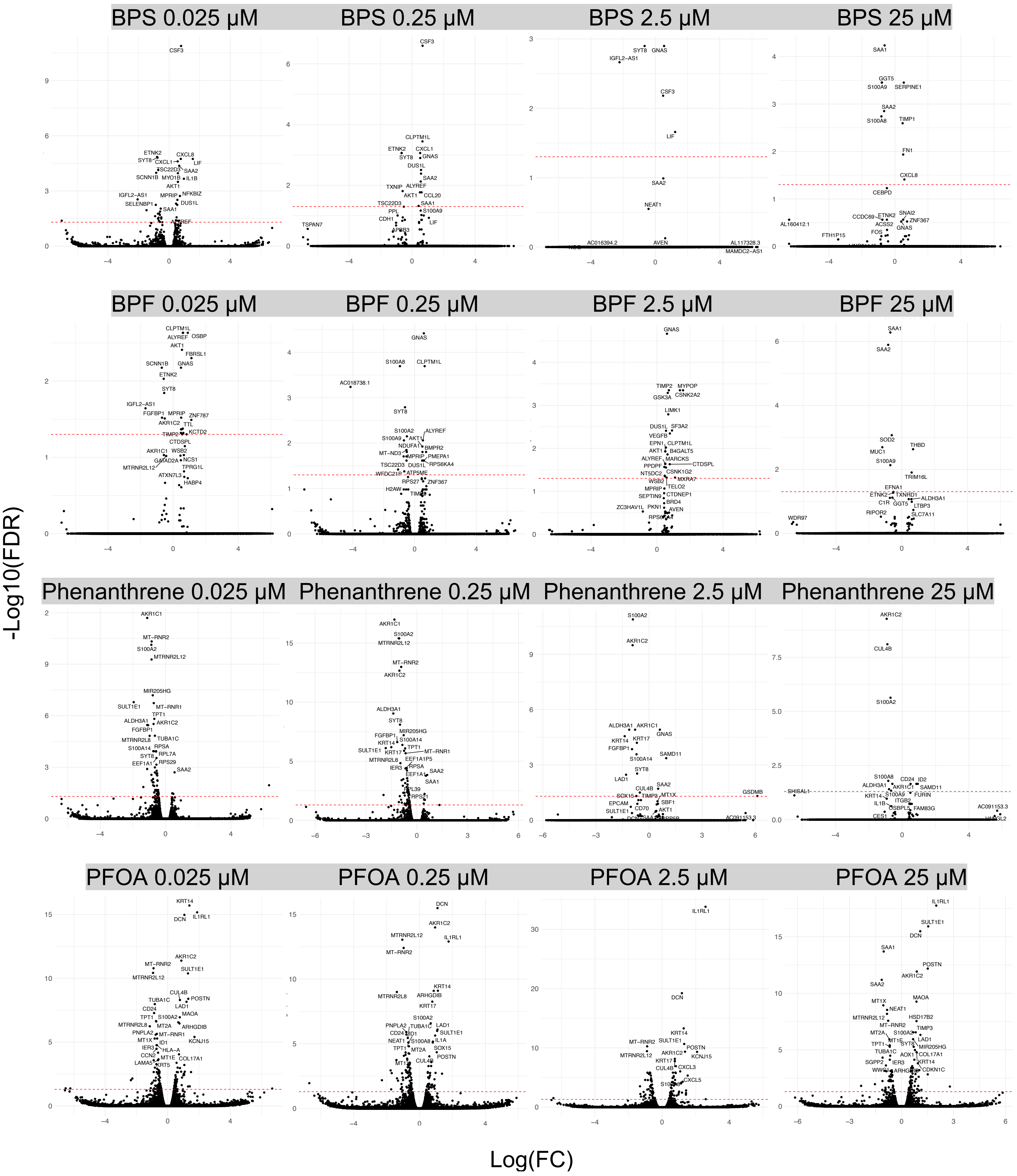

BPS 0.025 µM
BPS 0.25 µM
BPS 2.5 µM
BPS 25 µM
BPF 0.025 µM
BPF 0.25 µM
BPF 2.5 µM
BPF 25 µM
Phenanthrene 0.025 µM
Phenanthrene 0.25 µM
Phenanthrene 2.5 µM
Phenanthrene 25 µM
PFOA 0.025 µM
PFOA 0.25 µM
PFOA 2.5 µM
PFOA 25 µM
-Log10(FDR)
Log(FC)

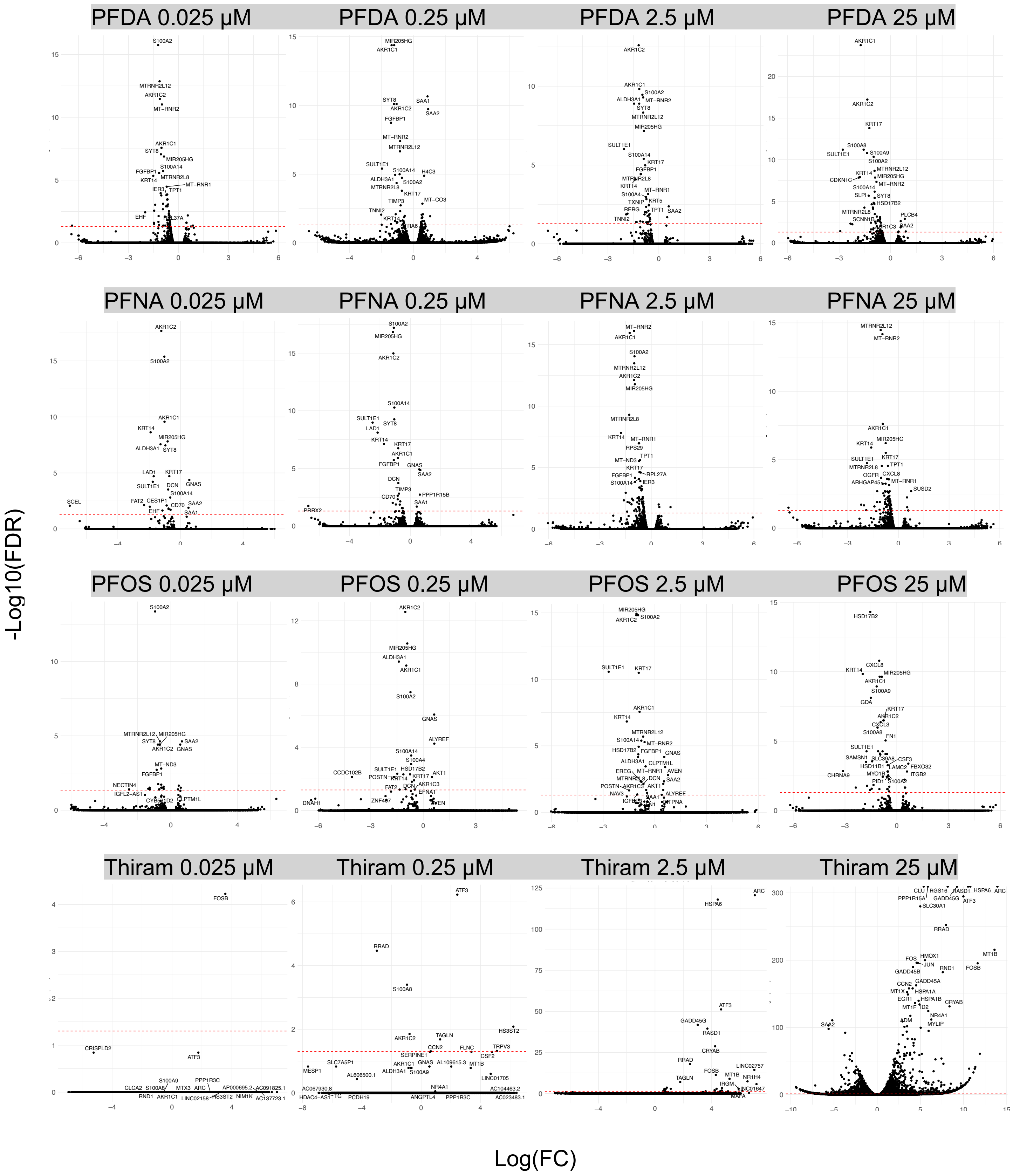
PFDA 0.025 µM
PFDA 0.25 µM
PFDA 2.5 µM
PFDA 25 µM
PFNA 0.025 µM
PFNA 0.25 µM
PFNA 2.5 µM
PFNA 25 µM
PFOS 0.025 µM
PFOS 0.25 µM
PFOS 2.5 µM
PFOS 25 µM
Thiram 0.025 µM
Thiram 0.25 µM
Thiram 2.5 µM
Thiram 25 µM
-Log10(FDR)
Log(FC)

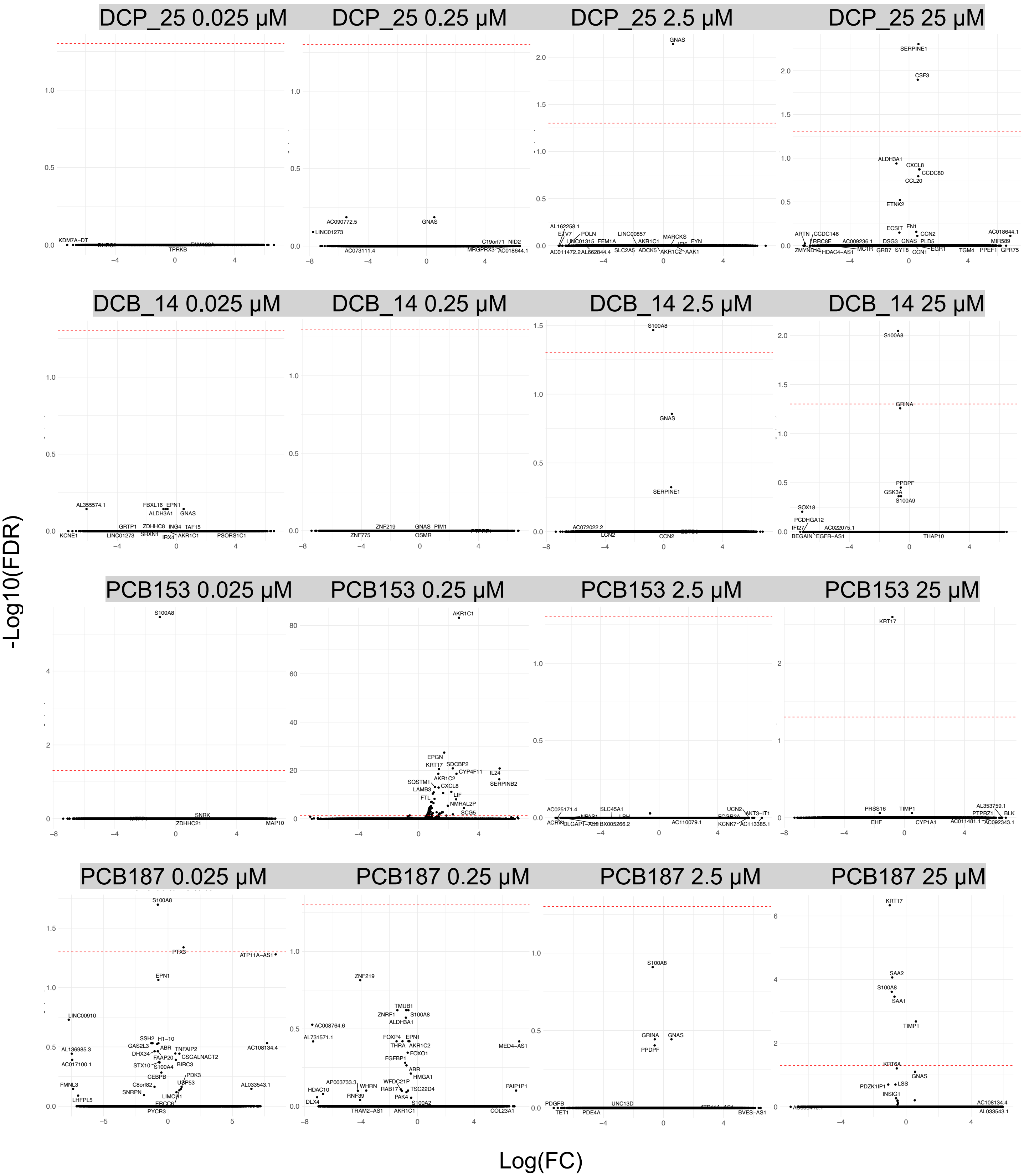
DCP_25 0.025 μM
DCP_25 0.25 μM
DCP_25 2.5 μM
DCP_25 25 μM
DCB_14 0.025 μM
DCB_14 0.25 μM
DCB_14 2.5 μM
DCB_14 25 μM
PCB153 0.025 μM
PCB153 0.25 μM
PCB153 2.5 μM
PCB153 25 μM
PCB187 0.025 μM
PCB187 0.25 μM
PCB187 2.5 μM
PCB187 25 μM
-Log10(FDR)
Log(FC)

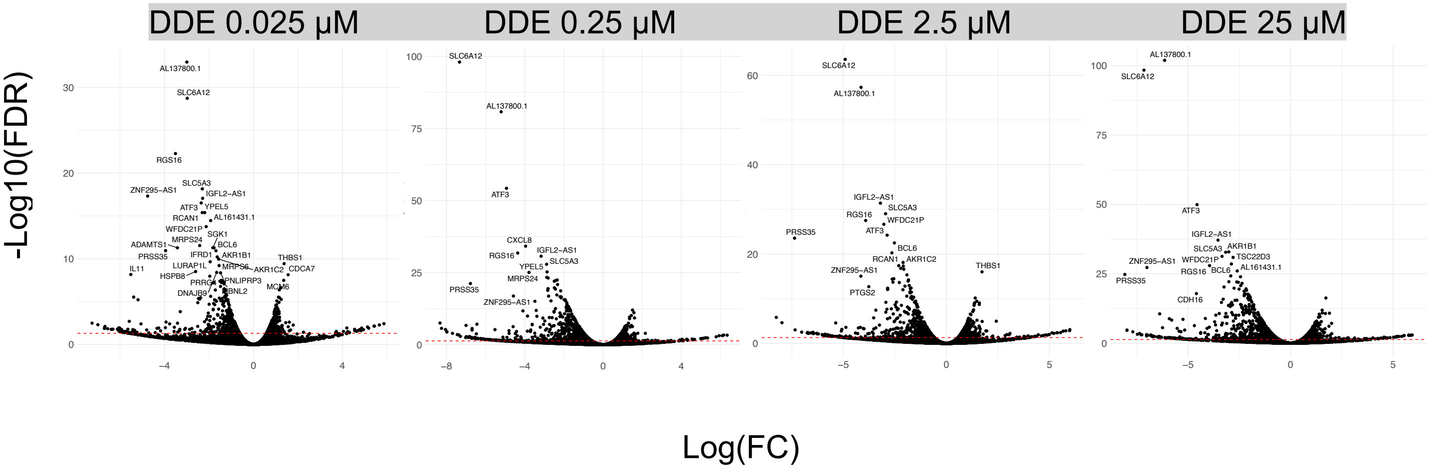


**Supplemental Figure 1: Differential gene expression for 21 disparities associated chemicals at all four chemical doses.** Differential gene expression between each chemical dose and controls was calculated using quasi-likelihood negative binomial generalized log-linear modeling in edgeR. Red lines mark a false discovery rate (FDR) adjusted p-value of 0.05.

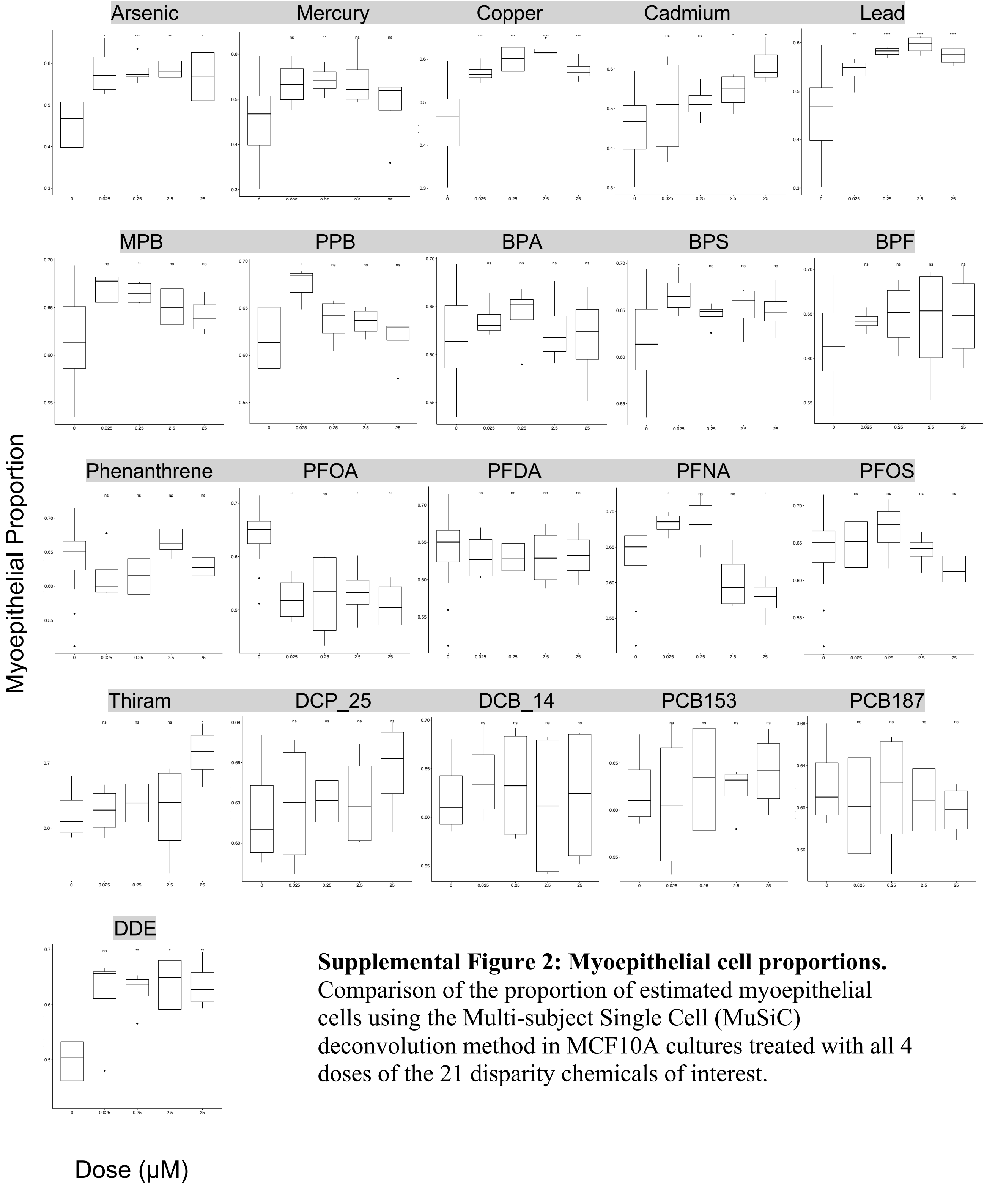


**Supplemental Figure 2: Myoepithelial cell proportions.** Comparison of the proportion of estimated myoepithelial cells using the Multi-subject Single Cell (MuSiC) deconvolution method in MCF10A cultures treated with all 4 doses of the 21 disparity chemicals of interest.

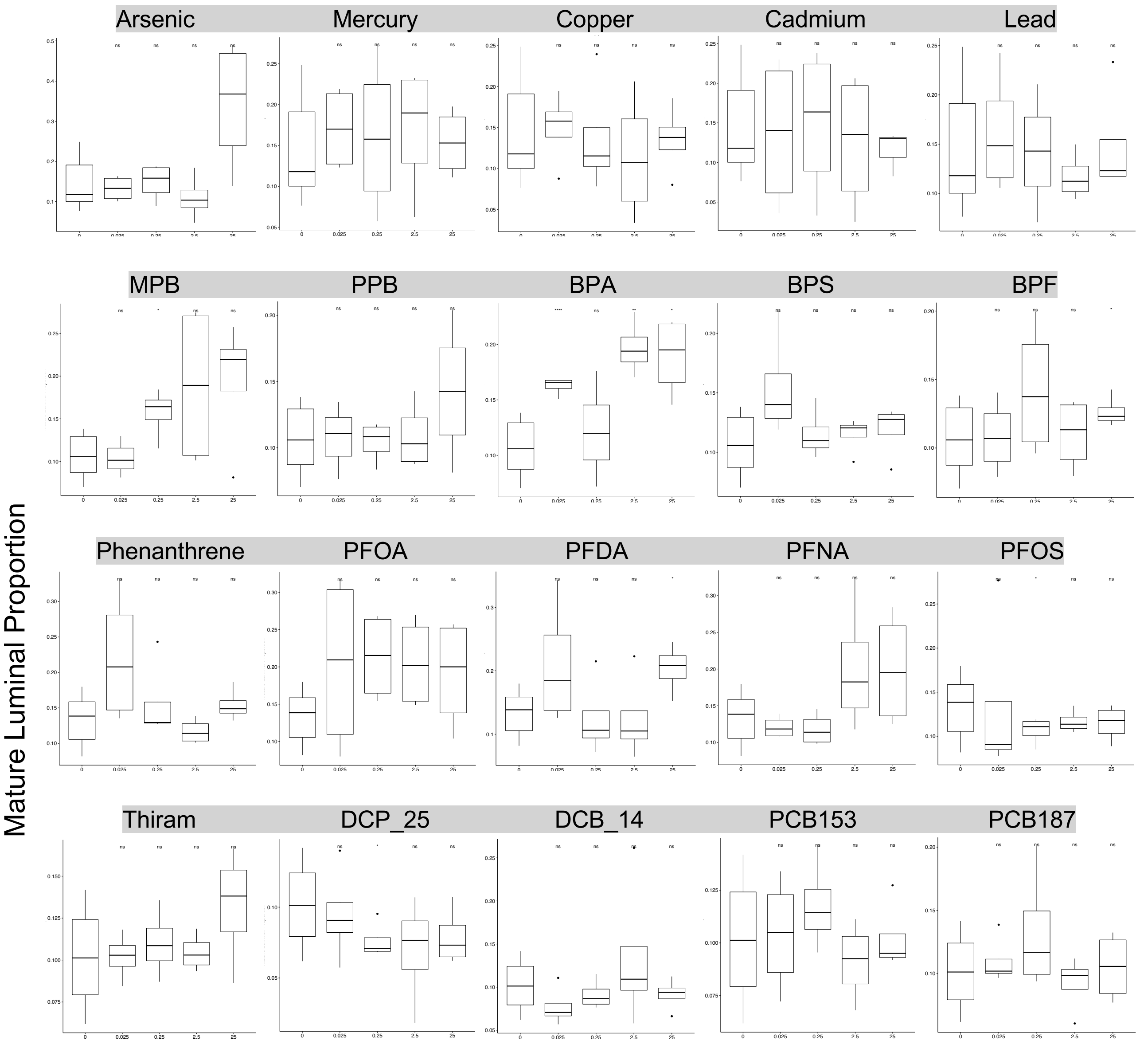


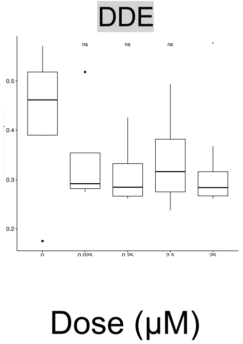


**Supplemental Figure 3: Mature luminal cell proportions.** Comparison of the proportion of estimated mature luminal cells using the Multi-subject Single Cell (MuSiC) deconvolution method in MCF10A cultures treated with all 4 doses of the 21 disparity chemicals of interest.

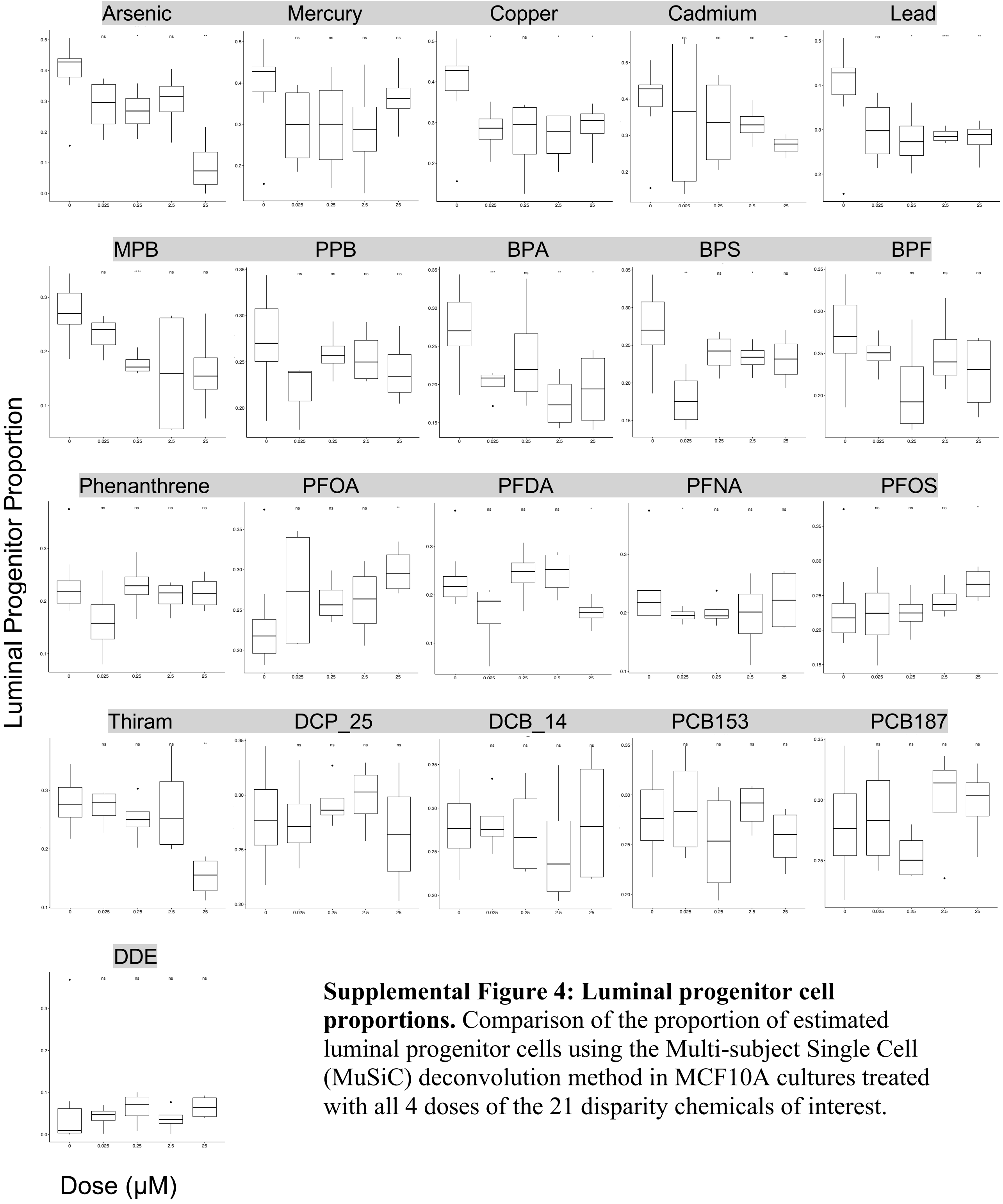


**Supplemental Figure 4: Luminal progenitor cell proportions.** Comparison of the proportion of estimated luminal progenitor cells using the Multi-subject Single Cell (MuSiC) deconvolution method in MCF10A cultures treated with all 4 doses of the 21 disparity chemicals of interest.
